# Auditory decision-making deficits after permanent noise-induced hearing loss

**DOI:** 10.1101/2024.09.23.614535

**Authors:** Madeline P. Berns, Genesis M. Nunez, Xingeng Zhang, Anindita Chavan, Klavdia Zemlianova, Todd M. Mowery, Justin D. Yao

**Author notes:** Correspondence: Justin D. Yao, Ph.D., Rutgers, The State University of New Jersey Nelson Biological Laboratory D418, 604 Allison Road, Piscataway, NJ 08854. Co-first Author (Madeline P. Berns, Genesis M. Nunez).

## Abstract

Loud noise exposure is one of the leading causes of permanent hearing loss. Individuals with noise-induced hearing loss (NIHL) suffer from speech comprehension deficits and experience impairments to cognitive functions such as attention and decision-making. Here, we tested whether a specific sensory deficit, NIHL, can directly impair auditory cognitive function. Gerbils were trained to perform an auditory decision-making task that involves discriminating between slow and fast presentation rates of amplitude-modulated (AM) noise. Decision-making task performance was assessed across pre-versus post-NIHL sessions within the same gerbils. A single exposure session (2 hours) to loud broadband noise (120 dB SPL) produced permanent NIHL with elevated threshold shifts in auditory brainstem responses (ABRs). Following NIHL, decision-making task performance was tested at sensation levels comparable to those prior to noise exposure in all animals. Our findings demonstrate NIHL diminished perceptual acuity, reduced attentional focus, altered choice bias, and slowed down evidence accumulation speed. Finally, video-tracking analysis of motor behavior during task performance demonstrates that NIHL can impact sensory-guided decision-based motor execution. Together, these results suggest that NIHL impairs the sensory, cognitive, and motor factors that support auditory decision-making.

## Introduction

Sensory deficits can impair cognitive function. For example, hearing loss reduces auditory processing skills and speech comprehension [1–4], and many forms of peripheral hearing loss can impair several domains of cognitive function that are independent of hearing sensitivity, like attentional focus and decision-making [5–8]. However, studies that directly assess the effects of noise-induced hearing loss (NIHL) on cognitive function are limited. Furthermore, it is difficult to distinguish whether NIHL-related cognitive and perceptual deficits arise from peripheral or central mechanisms.

Exposure to loud nose is the leading cause of acquired hearing loss [9]. The perceptual effects of NIHL can vary among individuals, and while NIHL-related damage to peripheral function may account for issues concerning auditory perception and speech recognition [10–11], inter-subject differences could be attributed to deficits in central or top-down processing mechanisms [12–13]. In fact, recent evidence suggests “non-sensory” cognitive factors may be vulnerable to NIHL. One study reported individuals with NIHL from occupational noise exposure exhibited poorer working memory, less attentional focus, and slower reaction times compared to normal-hearing counterparts [14]. These NIHL impairments can occur immediately after a period of loud noise exposure [15]. This raises the question of which specific sensory and non-sensory factors that support cognitive function are vulnerable to NIHL.

In the current study, we assessed auditory decision-making task performance across normal-hearing and NIHL conditions within the same animals. Auditory decision-making is a cognitive process that involves transforming accumulated sensory inputs into categories that guide perceptual judgement and action [16]. We tested the hypothesis that NIHL leads to cognitive deficits by impairing sensory and non-sensory factors that support auditory decision-making. First, we found that gerbils are capable of performing a sound-guided decision-making task that involves distinguishing between slow (<6.25-Hz) and fast (>6.25-Hz) amplitude-modulated (AM) broadband noise. Second, exposure to loud noise for one period of 2 hours permanently decreased hearing sensitivity. After NIHL, perceptual acuity and attentional focus were degraded, choice bias shifted, evidence accumulation speed slowed down, and motor function was disrupted. To control for differences in acoustic sensitivity across hearing status conditions, we adjusted the stimulus sound level to ensure auditory signals were presented at comparable sensation levels after NIHL. Altogether, we found that NIHL impairs the sensory, non-sensory, and cognitive factors supporting auditory decision-making.

## Results

### Exposure to loud noise induces permanent hearing loss

Adult gerbils (N = 7) were exposed to 120-dB SPL broadband noise during a single period of 2 hours to induce permanent hearing loss (Figure 1A). Figure 1B displays example auditory brainstem responses (ABRs), a general measure of brainstem function, from one gerbil recorded pre- (“Baseline”) and 14 days post-noise exposure. We found no significant difference between males (n = 4) and females (n = 3) for ABR thresholds for clicks across 14 days post-noise exposure (repeated measures two-way ANOVA, F(4,20) = 0.17, p = 0.95). Figure 1C displays ABR thresholds for clicks as a function of 0, 1-, 7-, 14-, and 21-days post noise exposure. We found that ABR thresholds for clicks significantly increased across 21 days post-noise exposure (repeated measures one-way ANOVA, F(4,24) = 78, p < 0.0001). Threshold shifts were permanent and remained stable after 14 days post-noise exposure (Figure 1D) (post-hoc multiple comparisons, p > 0.05; mean ± SEM shift after Day-14 post-noise exposure = 41.4 dB SPL ± 4.96), demonstrating permanent noise-induced hearing loss (NIHL). This validates the different sound levels used during behavioral testing across hearing status conditions to control for shifts in hearing sensitivity (normal hearing = 50 dB SPL; NIHL = 90 dB SPL).

**Figure 1.**
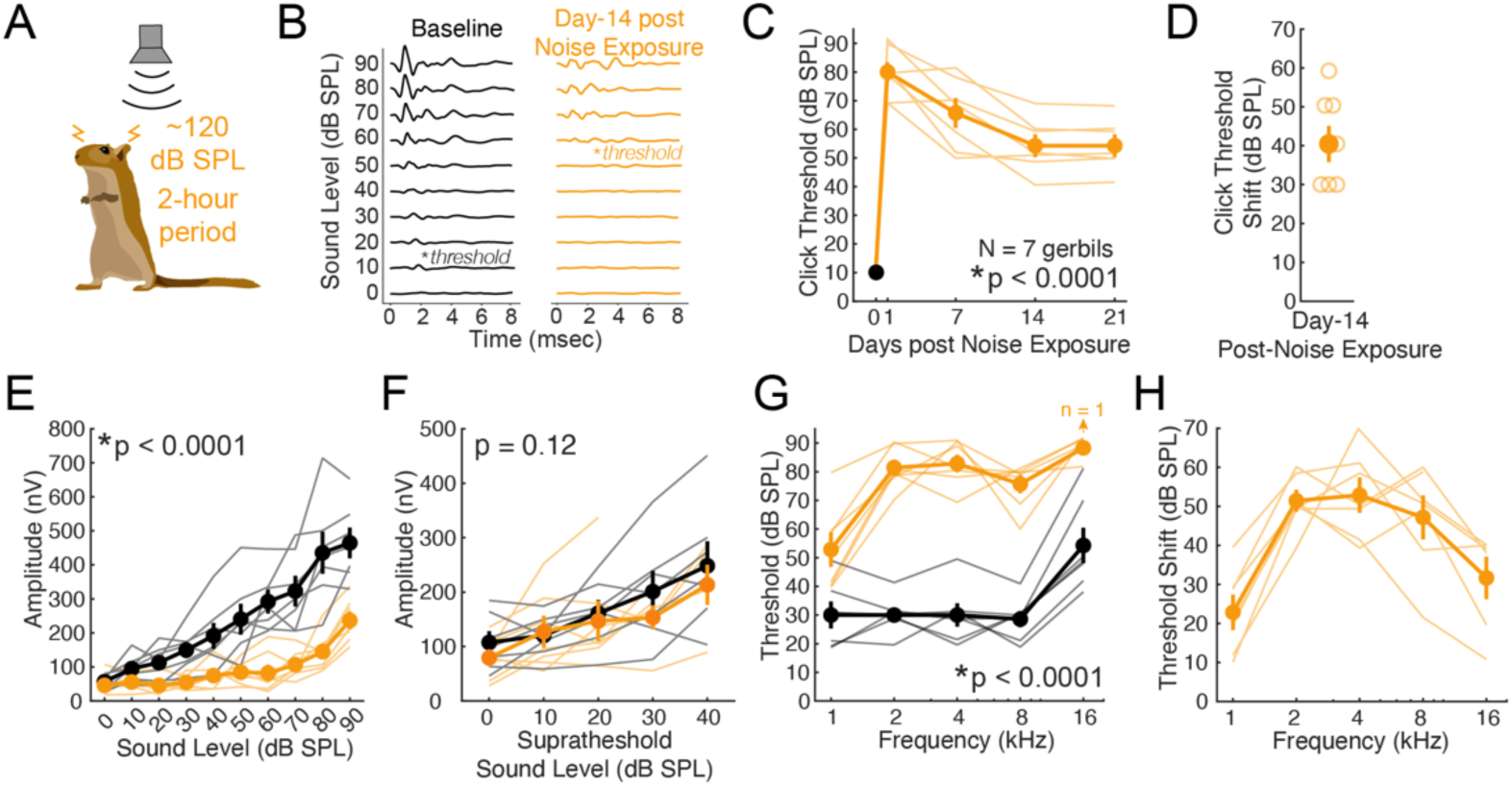
Exposure to loud noise leads to permanent hearing loss. (**A**) Gerbils are exposed to loud broadband noise (120 dB SPL) for one period of 2 hours. (**B**) Example ABR recordings to click stimuli during pre- (Baseline; black) and 14 days post-noise exposure (orange) sessions. ABR thresholds are determined as the lowest sound level to elicit a driven response (asterisk; Baseline: 10 dB SPL; Post-NIHL: 50 dB SPL). (**C**) Click threshold as a function of days (0, 1, 7, 14, and 21) post-noise exposure across all tested animals (N = 7). (**D**) Click threshold shift after 14 days noise-exposure for all animals. (**E**) Maximum ABR amplitudes for clicks as a function of sound level for pre- (black) versus 14 days post-noise exposure (orange). (**F**) Maximum ABR amplitudes for clicks as a function of sound level above each animal’s corresponding threshold for pre- (black) versus 14 days post-noise exposure (orange). (**G**) Thresholds for frequency tones across all tested animals for pre- (black) and 14 days post-noise exposure (orange). (**H**) Threshold shift as a function of frequency tones between pre- versus 14 days post-noise exposure for all tested animals. Symbols and error bars represent mean ± SEM.

We measured maximum amplitudes from click-evoked ABR responses as a function of sound level (dB SPL). Figure 1E displays results for all test animals for ABR recording sessions before (black) and 14 days after (orange) loud noise exposure. We found that maximum ABR amplitudes demonstrated a significant interaction between hearing status and sound level (repeated measures two-way ANOVA, F(9,120) = 5.85, p < 0.0001). For both pre- and post-NIHL conditions, maximum click-evoked ABR amplitudes significantly increased with louder sound levels (F(9,120) = 26.6, p < 0.0001). However, maximum click-evoked ABR amplitudes were significantly lower following NIHL compared to normal-hearing pre-noise exposure conditions (F(1,120) = 141.3, p < 0.0001). The differences in evoked ABR amplitudes between hearing status conditions could be due to differences in hearing sensitivity (i.e., increased threshold after NIHL). Thus, we compared maximum click-evoked ABR amplitudes across hearing status conditions as a function of suprathreshold sound levels (Figure 1F). Maximum ABR amplitudes grew significantly larger as suprathreshold sound levels increased (repeated measures two-way ANOVA, F(4,60) = 6.07, p < 0.0001), but there was no main effect of hearing status (repeated measures two-way ANOVA, F(1,4) = 2.44, p = 0.12), nor was there an interaction between hearing status and suprathreshold sound level (repeated measures two-way ANOVA, F(4,60) = 0.45, p = 0.77).

We also measured ABR thresholds across frequency tones (1, 2, 4, 8, and 16-kHz) (Figure 1G). ABR thresholds for tone frequencies across 21 days post-noise exposure followed a similar trend to ABR thresholds for clicks. For each tested tone frequency, ABR thresholds permanently increased across 21 days post-noise exposure (repeated measures one-way ANOVA (F(4,24) = 17.7-60.9, p < 0.0001) and remained stable after 14 days post-noise exposure (post-hoc multiple comparisons, p > 0.05). After 14 days post-noise exposure, ABR thresholds displayed a significant interaction between hearing status (pre- versus post-NIHL) and tone frequencies (repeated measures two-way ANOVA, F(4,60) = 6.06, p < 0.0001). In addition, we found a significant main effect of hearing status, with ABR thresholds across all tested frequencies displaying a significant increase 14 days post-noise exposure (two-way ANOVA, F(1,60) = 321.6, p < 0.0001). NIHL- related shifts in ABR thresholds were more prominent for 2-, 4-, and 8-kHz (Figure 1H).

### Permanent noise-induced hearing loss impairs auditory decision-making

To determine whether permanent NIHL impairs auditory decision-making, we trained adult gerbils with normal hearing (N = 7) to perform a single-interval alternative forced choice (AFC) AM rate discrimination task (Figure 2). Briefly, gerbils are trained to self-initiate trials by placing their nose in a nose port, after which they discriminate between slow (<6.25-Hz) versus fast (>6.25-Hz) amplitude-modulated (AM) broadband noise rates by approaching the left or right food trough, respectively. Psychometric testing after NIHL was assessed 14 days post-noise exposure since this is the timepoint when ABR thresholds had permanently shifted and stabilized. To control for differences in overall hearing sensitivity after NIHL, we adjusted the sound level of presented acoustic stimuli during behavioral sessions following NIHL. For behavioral testing under normal-hearing conditions, acoustic stimuli were presented at a sound level of 50 dB SPL. After NIHL, acoustic stimuli were presented at a sound level of 90 dB SPL. Across both hearing status conditions, acoustic stimuli were presented at a similar sound level above threshold (Wilcoxon signed-rank test, p = 1; sensation level pre-NIHL: median = 40 dB SPL above threshold; sensation level post-NIHL: median = 40 dB SPL above threshold). Thus, any differences in behavioral performance could not be attributed to poorer auditory sensitivity after NIHL because behavioral testing was performed at similar sensation levels.

**Figure 2.**
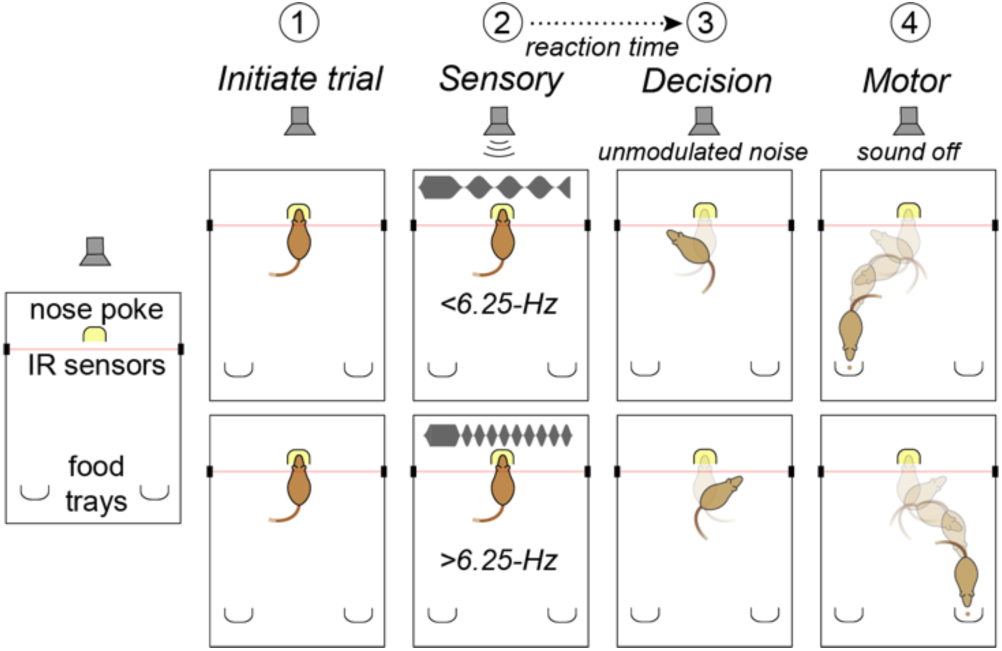
Schematic of the single-interval, two-AFC auditory decision-making task. (1) Gerbils self-initiate trials by placing their nose in a nose port. (2) AM signal turns on. (3) Gerbils deliberate decision and then leave nose port area that is designated by IR sensors. Once gerbils leave this area the AM signal transitions into unmodulated noise. (4) Gerbils approach the left or right food trough based on decision (<6.25-Hz or >6.25-Hz).

Psychometric functions for each animal across all pre- (black) and post-NIHL (orange) sessions are displayed in Supplemental Figure 1. We fit psychometric functions to a gaussian distribution and compared the fitted parameters of Threshold and Slope across hearing status conditions. Slope represents the degree of perceptual acuity and Threshold represents choice bias relative to the “choose left versus right” boundary of 6.25-Hz. Following NIHL, all animals displayed a significant decrease in psychometric Slope (two-way t-test, t = -10.8 to -1.89, p < 0.0001) and a significant increase in Lapse Rate (two-way t-test, t = -13.2 to -6.77, p < 0.0001). Five out of the seven animals displayed a significant shift in psychometric Threshold after NIHL (two-way t-test, t = -15.2 to -2.33, p < 0.05).

To further assess auditory decision-making ability across hearing status conditions, we calculated psychometric Slope, Threshold, and Lapse Rate values for each animal by pooling task performance across all corresponding sessions. Figure 3A displays a psychometric function from one example animal across pre- (black) and post-NIHL (orange) sessions. Auditory decision-making task performance was significantly impaired after NIHL, with animals displaying a significant decrease in Slope (two-way t-test, t = 24.2, p < 0.0001; Figure 3B), a significant shift in Threshold (two-way t-test, t = 5.29, p = 0.006; Figure 3C, D), and a significant increase in Lapse Rate (two-way t-test, t = -16.1, p < 0.0001; Figure 3E). These results suggest that NIHL impacts auditory decision-making ability.

**Figure 3.**
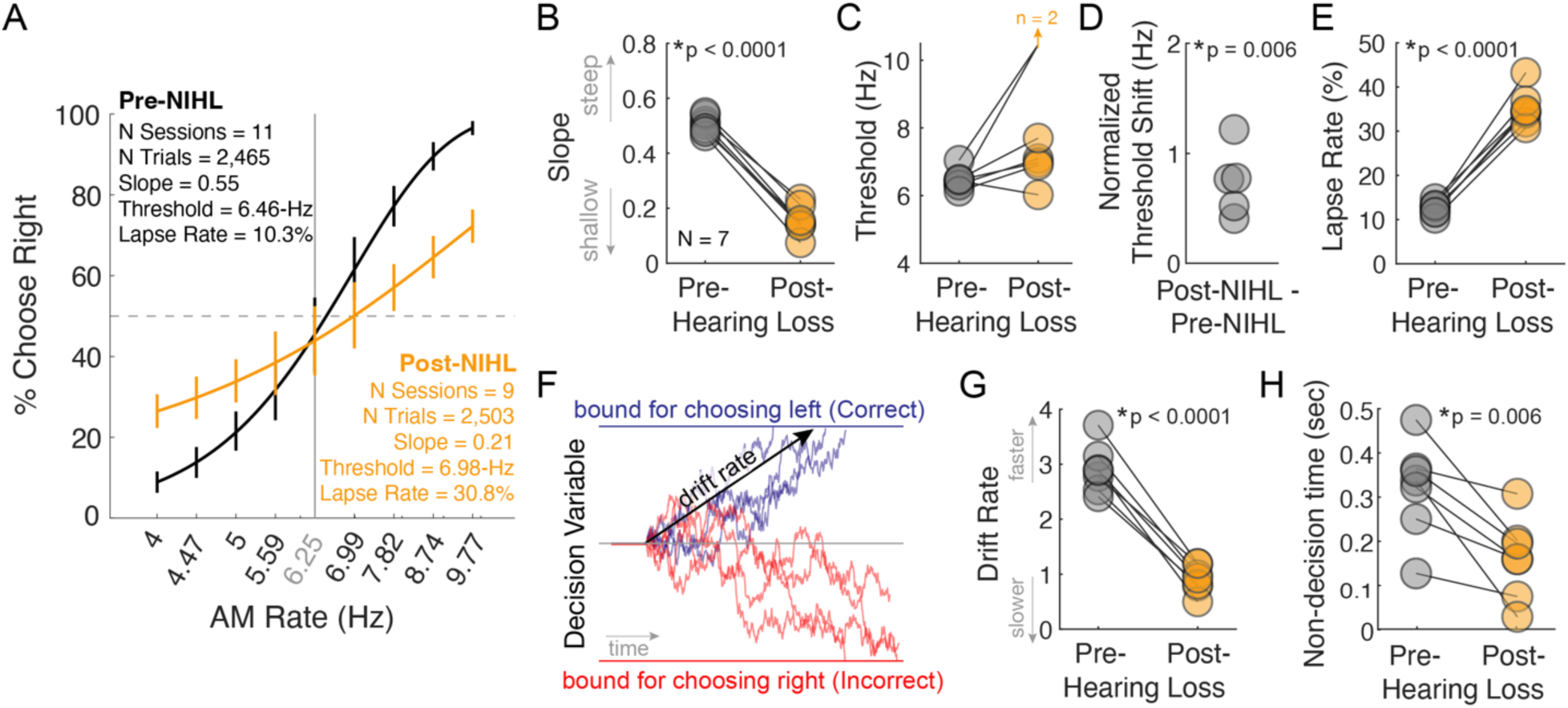
NIHL impairs auditory decision-making. (**A**) Example psychometric performance from one animal for Pre- (black) versus Post-NIHL (orange) conditions. Psychometric functions were constructed from collapsing all trials from each test session. Error bars represent 95% confidence intervals. (**B**) Corresponding psychometric Slope values across Pre- versus Post-NIHL for all animals. (**C**) Corresponding psychometric Threshold values across Pre- versus Post-NIHL for all animals. 2 animals did not possess a Threshold value after NIHL. (**D**) Normalized shift in psychometric Thresholds after NIHL. (**E**) Corresponding psychometric Lapse Rate values across Pre- versus Post-NIHL for all animals. (**F**) Schematic of DDM decision-making process for several trials across AM rates (AM rate <6.25-Hz; Correct = choose left). (**G**) Corresponding Drift Rate values across Pre- versus Post-NIHL for all animals. (**H**) Corresponding Non-decision time values across Pre- versus Post-NIHL for all animals. P-values represent statistical outcomes from paired- sample t-tests.

We assessed the impact of NIHL on the cognitive process of evidence accumulation by fitting task performance to a drift diffusion model (DDM). This allowed us to quantify decision-making variables such as “Drift Rate”. Figure 3F displays a schematic of the DDM decision-making process for several trials. Drift Rate is the average slope of the decision variable across time during a given trial and is a highly cognitive process supporting sensory-guided decision-making. Drift Rate represents the speed of evidence accumulation, where larger Drift Rate values exemplify faster evidence accumulation processing. After NIHL, Drift Rates significantly decreased (two-way t-test, t = 13.9, p < 0.0001; Figure 3G), demonstrating diminished speed of acoustic evidence accumulation during the decision-making process. NIHL did not significantly alter a separate DDM parameter of “Boundary separation” (two-way t-test, p = 0.22). However, an additional DDM parameter of “Non-decision time”, which reflects sensory encoding and motor execution, was significantly reduced after NIHL (two-way t-test, t = 4.12, p = 0.006; Figure 3H). This is similarly reflected by NIHL-related impairments to perceptual acuity (Figure 3A, B) and reaction times (RT; see below). We found no significant difference between males (n = 4) and females (n = 3) for NIHL-attributable shifts in task performance (Wilcoxon rank sum test: Slope p = 0.86, Threshold p = 0.40, Lapse Rate p = 0.11, Drift Rate p = 0.40, Non-decision time p = 0.63). Slope and Threshold measures from psychometric performance represent sensory processing factors. Lapse Rate represents a non-sensory factor, and Drift Rate and Non-decision time measures may serve as proxies for cognitive processing factors during decision-making. Thus, our results demonstrate NIHL impairs sensory, non-sensory, and cognitive processing abilities of auditory decision-making.

### Noise-induced hearing loss effects on motor function

Since animals displayed impairments to auditory decision-making task performance following NIHL, we examined whether NIHL impacts motor function. We first compared reaction times (RTs) for correct trials as a function of AM rates between hearing status conditions. We found that RTs as a function of AM rates were similar for 6/7 animals across pre- and post-NIHL conditions (two-way ANOVA, F(1,14-21) = 1.04-2.79, p = 0.05-0.47) (Supplemental Figure 1). When compared RTs across trial type (Correct versus Incorrect trials) and hearing status (Pre- versus Post-NIHL), we observed that 6/7 animals displayed significant differences in RTs between trial type (two-way mixed model ANOVA, F(1,4610 – 5897) = 8.82 – 56.1, p < 0.0001) and hearing status (two-way mixed model ANOVA, F(1,4610 – 5897) = 6.49 – 96.9, p < 0.0001) (Supplementary Figure 2). Five animals exhibited faster RTs for incorrect trials during normal-hearing conditions, which could be attributed to a lack of attentional control for appropriately processing task-related auditory signals. This potential lack of attentional control is further exhibited after NIHL, where RTs were typically faster than their normal-hearing counterparts for both correct and incorrect trials (Supplementary Figure 2).

To further characterize potential differences in motor function before and after NIHL, we analyzed video tracking data from all recorded videos during task performance. We specifically tracked each animal’s body movements (i.e., head, torso, and tail position) with the pose-tracking algorithm SLEAP [17] (Figure 4A, B). We found that the durations of decision-based motor execution (i.e., the duration of a completed full turn towards the left or right food trough) varied across individual animals but were significantly affected by NIHL in 6/7 animals (two-way mixed model ANOVA, F(1,4380-6209) = 93.1-886.6, p < 0.0001) (Figure 4C). Notably, after NIHL, decision-based motor execution durations increased significantly in two of three female animals (two-way mixed model ANOVA, F(1,5075-6209) = 46.4-271.7, p < 0.0001) and decreased significantly in all four males (two-way mixed model ANOVA, F(1,4380-4931) = 93.1-886.6, p < 0.0001). Thus, decision-based motor execution was significantly impacted after NIHL. In addition, our results demonstrate potential sex-related differences to motor function after NIHL with females exhibiting slower decision-based motor execution than males.

**Figure 4.**
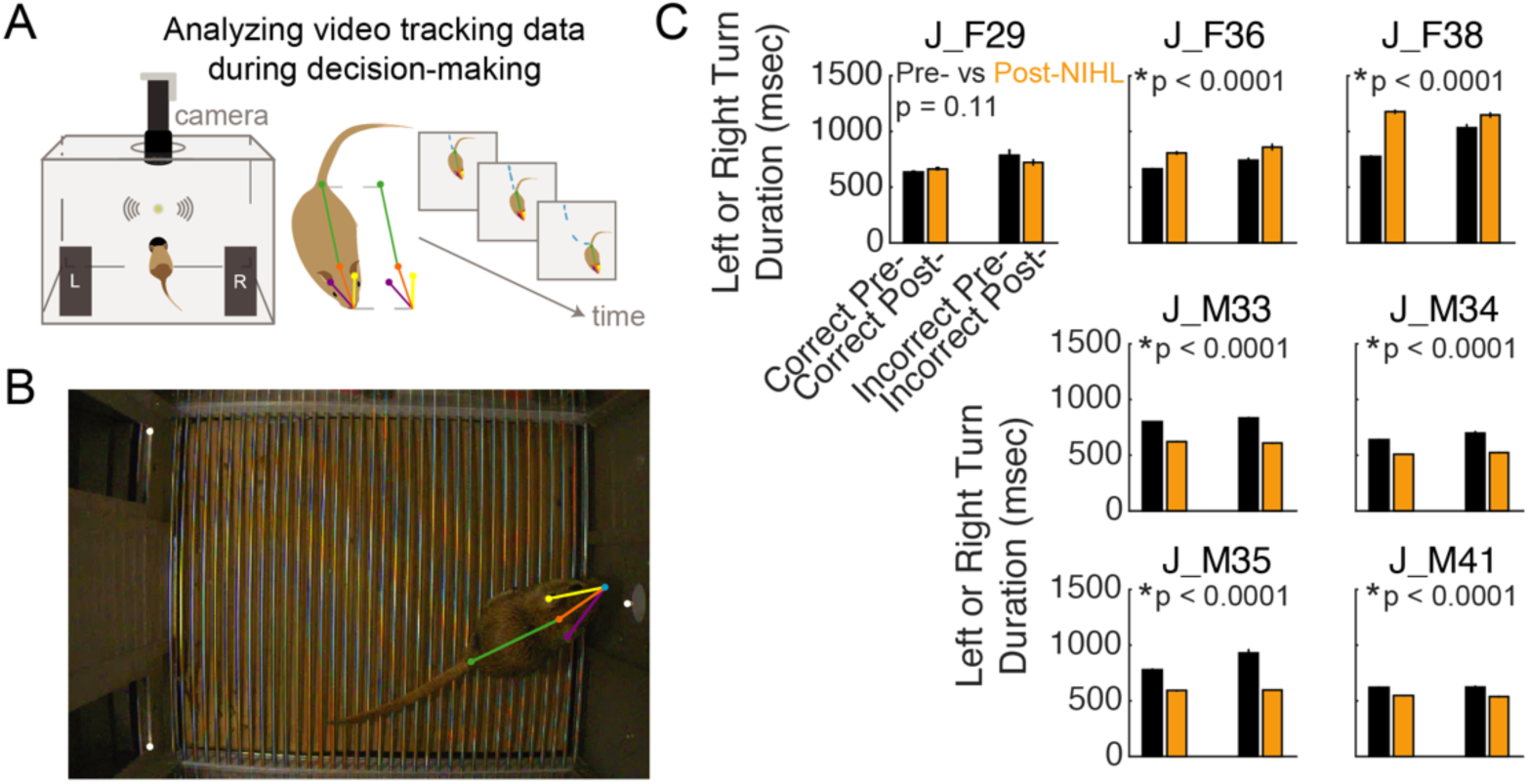
Video tracking analyses across Pre- versus Post-NIHL conditions. (**A**) Task sessions are video recorded and pose-tracking is performed with SLEAP [17]. (**B**) Example video frame showing behavioral testing arena. Nose port and food troughs locations are marked by white dots. Tracking of the gerbil’s nose, head, ears, and tail are indicated by the colored skeleton. (**C**) Mean ± SEM turn duration grouped across different trial outcomes (Correct and Incorrect) and hearing status (Pre- and Post-NIHL) for each animal. J_F29, J_F36, and J_F38 are the IDs assigned to female gerbils. J_M33, J_M34, J_M35, and J_M41 are the IDs assigned to male gerbils. Asterisks denote a statistically significant interaction of trial outcome and hearing status on turn duration (two-way mixed model ANOVA). For a subset of the data, error bars are small and lie within the mean data symbol.

## Discussion

Auditory decision-making is a cognitive process that relies on sensory and non-sensory factors. Here, we asked whether NIHL impairs the discrimination of time-varying acoustic cues during auditory decision-making in gerbils. We demonstrated that a single exposure period of 2 hours to very loud noise (120 dB SPL) induces permanent hearing loss. After assessing auditory decision-making before and after noise-exposure, we found that NIHL impairs sensory, non-sensory, and cognitive factors that support sound-guided decision-making. Taken together, our findings suggest that a specific sensory deficit, NIHL, leads to detrimental effects on cognitive function.

Loud noise exposure primarily damages auditory peripheral structures and can have long-lasting effects on auditory function [11]. The most common outcome of loud noise exposure is elevated audiometric thresholds, or difficulty hearing sounds. Noise injury often includes loss of synapses between inner hair cells and auditory nerve fibers [11, 18]. This leads to degraded transmission of acoustic signals along the ascending auditory pathway to auditory cortex, resulting in deficits to auditory perceptual skill [19–20]. Our results demonstrate that in gerbils, peripheral damage via loud noise exposure (Figure 1) produces auditory perceptual and cognitive deficits even when sensation level is normalized across hearing status conditions (Figure 3). This could be attributed, in part, to functional changes along the ascending auditory system originating from cochlear damage [10–11, 21–24].

Many forms of peripheral hearing loss can impair several domains of cognitive function, including working memory, attentional focus, and decision-making, which can be independent of hearing sensitivity [5–8, 24–25]. This is supported by several studies demonstrating learning deficiencies after auditory deprivation [26], and deficits to auditory and visual task performance after induced dysfunction of the inner ear [27]. In addition, NIHL presents an increased risk for cognitive impairment exhibited through deficits in spatial learning and memory [28–30] and temporal order object recognition tasks [31]. We extend these findings in the context of cognitive assessment through sound-guided decision-making. The process of decision-making involves cognitive components that can be measured and compared between clinical conditions. We found that NIHL can disrupt sensory-guided motor execution and provide evidence suggesting sex-related differences in how NIHL impacts motor function (Figure 4C). This suggests that NIHL can interfere with cognitive-motor interactions, thus broadening the reach of NIHL-related effects across multiple sensory, cognitive, and motor domains. It is possible that NIHL-related effects to motor function are due to its impact on balance and vestibular function, which may be attributed to inflammation within the cochlea [32–33]. Alternatively, NIHL may impair the corollary discharge signals between auditory and motor cortices that support sound-guided behavior [34]. Overall, we found that NIHL can detrimentally impact the sensory and non-sensory factors that support auditory-guided decision-making. This has strong significance for assessing the impact of hearing loss on cognitive decline, as individuals with hearing loss are at greater risk for incident cognitive impairment and Alzheimer’s-related dementias [35–36].

NIHL-related impairments to cognitive function could be attributed to deficits along the auditory pathway and its downstream targets [37–39]. In fact, there is some evidence that individuals with hearing loss exhibit reduced attentional modulation of cortical responses, leading to poor performance on selective attention tasks [40–41]. Furthermore, several reports propose that functional connectivity between central auditory areas and higher cortical regions involved in cognitive function are disrupted after hearing loss [42–45]. In rodents, NIHL initiates plasticity downstream of auditory cortex that alters neural responses in the amygdala and striatum [46], and changes functional connectivity between auditory and non-auditory cortical regions, such as frontal cortex, primary motor cortex, cingulate cortex, hippocampus, and cerebellum [47–50]. Disruptions in functional connectivity between visual cortex and frontal cortex after NIHL are also reported [47], with shifts along functional boundaries within audiovisual cortex [51]. Overall, deprivation-induced plasticity beyond the auditory central pathway may explain cognitive function impairments that accompany hearing loss.

Hearing loss studies typically focus on outcomes regarding auditory sensation and perceptual capabilities. However, it is robustly evident that hearing loss can lead to cognitive dysfunction, whether it be guided by sensory and/or non-sensory factors. Here, we found that, even when controlling for loss of auditory sensitivity, NIHL leads to auditory cognitive deficits by impairing the sensory and non-sensory factors that guide sound-driven decision-making. One plausible hypothesis that explains our results is that NIHL degrades sensory and non-sensory processing mechanisms within and beyond the auditory cortical pathways. Previous reports demonstrate the pathway between auditory cortex to parietal cortex is involved in the transformation of encoded auditory cues for guiding sound-driven perceptual decisions [52–53]. Our future work will explicitly test whether specific candidate cortical circuits, such as auditory to parietal cortex, provide a neural explanation for hearing loss obstructions to cognitive function.

## Methods

### Experimental animals

Mongolian gerbils (*Meriones unguiculatus,* N = 7, 4 males) were weaned from commercial breeding pairs (Charles River) and housed on a 12-hr light/dark cycle with ad libitum access to food and water. All procedures were approved by the Institutional Animal Care and Use Committee at Rutgers University. Each experiment was performed once with technical replication occurring for behavioral data (i.e., each animal was tested psychometrically multiple times).

### Psychophysical testing

We assessed auditory decision-making in adult gerbils with a single-interval alternative-forced choice (AFC) appetitive conditioning paradigm. Adult gerbils were placed on controlled food access and trained to discriminate amplitude-modulated (AM) broadband noise (0.1-20 kHz; 100% modulation depth) presented at slow (<6.25-Hz) versus fast (>6.25-Hz) rates (Figure 2). Gerbil auditory decision-making task performance is conducted in a behavioral arena test cage (Med Associates) housed inside a sound-attenuating cubicle (Med Associates) or an sound attenuation booth (Whisper Room). Gerbils self-initiated trials by placing their nose in a nose poke port for a minimum of 100 msec that interrupted an infrared beam and triggered an acoustic stimulus. Each AM stimulus was presented at a sound pressure level (SPL) of 50 dB (before noise-induced hearing loss) or 90 dB (after noise-induced hearing loss) and had a 100 msec onset ramp, followed by an unmodulated period of 100 msec that transitioned to an AM signal. During acoustic stimulation, a gerbil approaches the left or right food trough. Infrared sensors placed around the nose poke area detect when gerbils make their initial approach to one of the two food troughs, and the stimulus immediately transitions to unmodulated noise. We calculate reaction time as the duration from AM signal onset to when animals departed the nose poke area. When the infrared beam at the correct food trough (<6.25-Hz = approach left tray; >6.25-Hz = approach right tray) is broken, a pellet dispenser (Med Associates) delivers one dustless precision pellet (20 mg; Bio-Serv). All task sessions are observed via a closed-circuit monitor. Stimuli, food reward delivery, and behavioral data acquisition were controlled by an iPac computer system running iCon behavioral interfaces (Tucker-Davis Technologies). Auditory stimuli were presented from a calibrated multifield speaker (MF1, Tucker-Davis Technologies) positioned 4 cm above the nose poke port. Sound calibration measurements were verified with a digital sound meter (Larson Davis SoundExpert 821 ENV).

AM stimuli consisted of 11 presentation rates in steps of equal logarithmic spacing: 4-, 4.47-, 5-, 5.59-, 6.25-, 6.99-, 7.82-, 8.74-, and 9.77-Hz. AM stimuli were randomly presented with the following probabilities: 35.3% = 4- and 9.77-Hz; 23.5% = 4.47- and 8.74-Hz; 23.5% = 5- and 7.82-Hz; 11.8% = 5.59- and 6.99-Hz; 5.9% = 6.25-Hz. This probability distribution is to ensure greater numbers of trials for “easy” AM rates and promote task engagement. The AM stimulus presented at 6.25-Hz represented a “catch” trial. During catch trials, a reward pellet is dispensed 50% of the time, regardless of the food trough the gerbil approaches. We chose to utilize AM stimuli for two main reasons: (1) AM stimuli are typically used to study the envelope information that is a feature of all natural sounds [54–56]; (2) The time-varying nature of AM stimuli enable the assessment of evidence accumulation during decision-making.

### Auditory brainstem response recording

Auditory brainstem response (ABR) recordings were conducted before noise-induced hearing loss, and 1-, 7-, and 14-days after noise exposure. Gerbils are anesthetized with isoflurane (1.0%) and placed in a small acoustic chamber (IAC, Sound Room Solutions). Pin electrodes were inserted subcutaneously at the vertex of the skull (positive electrode), caudal to the left pinna (inverting electrode), and into the left leg (ground). Stimulus generation, presentation, and data acquisition were conducted with Tucker-Davis Technologies’ BioSigRZ software and ABR system. Sound stimuli were delivered through a multi- field speaker (MF1, Tucker-Davis Technologies) with a 10-cm tube (closed field) inserted into the left ear and placed at the opening of the ear canal. Acoustic stimuli consisted of 100 µs clicks (90 to 0 dB SPL, 500 repetitions across 10-dB SPL steps) and 5 msec tones (2 msec linear ramp) at 1, 2, 4, 8, and 16 kHz (90 to 20 dB SPL, 500 repetitions across 10-dB SPL steps). Thresholds for each acoustic stimulus were measured as the lowest dB SPL that elicited a stimulus-evoked ABR.

### Noise-induced hearing loss (NIHL) model

Once gerbils performed a minimum of 10 psychometric testing sessions (>2,000 trials), they are exposed to loud noise to induce permanent hearing loss. Awake gerbils are placed in a single-housed cage devoid of bedding within a sound-attenuating cubicle (Med Associates) located inside a sound attenuation booth (WhisperRoom). Gerbils are exposed to broadband noise (0.5-20 kHz) at 120 dB SPL presented from an overhead speaker 13” above the single-housed cages for a single 2-hour period.

### Performance measures and statistical analyses

We fit psychometric curves to each animal’s choice data across all presented AM rates for each session, and across all sessions. Psychometric functions were generated by fitting choice data to a cumulative Gaussian function. Gaussian model coefficients were fit with Matlab’s fitlm function. “Threshold” and “Slope” values were generated by the gaussian model fits. Threshold values represent choice bias relative to the boundary of 6.25-Hz. Threshold values >6.25-Hz represents a bias towards rightward choice (i.e., reporting “fast” AM rates). Threshold values <6.25-Hz represents a bias towards leftward choice (i.e., reporting “slow” AM rates). Slope values represent the degree of perceptual acuity. Larger slope values represent superior discrimination performance. Lapse rate values are calculated by the average percentage of incorrect trials for the easiest auditory signals of 4- and 9.77-Hz.

We measure cognitive processing aspects of auditory decision-making task performance by fitting psychophysical and chronometric data (i.e., percentage of correct trials and reaction times) to a drift diffusion model (DDM) [57–59]. We use the DDM to measure the accumulation of auditory evidence during the AFC auditory decision-making task. In the DDM, signed (+/-) momentary sensory evidence accumulates over time to create a decision variable (e.g., choose left or right). This process continues until the decision variable reaches either an upper or lower bound, and the bound dictates the choice. We fit the DDM to individual gerbils’ behavior using PyDDM [60], through which we applied a maximum-likelihood procedure. Specifically, a constant weighting function is applied to the moment-by-moment sensory evidence (i.e., AM rate). We incorporated the same data utilized for comparing psychometric curves and parameters (Supplemental Figure 1).

This version of the model has 4 free parameters: Boundary Separation (*a*), Drift Rate (*d*), Starting Point *(z)* and Non-decision time *(Ter)*. The starting point (*z*), which represents a response bias, was kept constant at 0, generating a model that did not account for bias. Drift Rate *(d)* is the average slope of the evidence accumulation process across many trials across sessions. For each trial, *d* was scaled linearly with the logarithmic distance of that trial’s AM rate from the midpoint rate (6.25-Hz), thus scaling it with the difficulty of the task. Boundary Separation (*a*), or the difference between the two available choices (i.e. left or right), is a measure of response caution. Non-decision time (*Ter*) is comprised of the time required to encode the stimulus and execute a motor action [61–62].

For each trial, the parameters were fitted according to the following equation that represents a standard DDM:

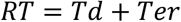

where *Td* represents the time required to make a decision, as calculated by the drift rate multiplied by the decision variable [62].

### Video recordings

We recorded video of each animal’s behavioral test session with a USB camera at 60 frames per second (Basler ace2). We specifically tracked the position of the gerbil’s nose, head, left and right ear, and tail base using the SLEAP algorithm [17]. We generated a trained gerbil network that consisted of >10,000 iterations and labeled frames. A gerbil’s angular velocity was calculated as the displacement of the head-to-tail vector over time. To reduce noise, we applied a moving average filter to smooth the raw angular data. Angular turn duration was then calculated as the time between stimulus onset and the moment at which the angular change of the gerbil plateaued, as identified via sustained edge detection logic.

Statistical analyses and procedures were implemented with custom-written Matlab scripts that incorporated the Matlab Statistics Toolbox, or custom-written Python scripts that incorporated modules with mathematical statistics functions, including SciPy and NumPy. For normally- distributed data (assessed by the Lilliefors test), we report mean ± SEM unless otherwise stated. For post-hoc multiple comparisons analyses, alpha values were Bonferroni-corrected. We used non-parametric statistical tests when data are not normally distributed.

## Acknowledgements

This work was supported by R00DC018600 (JDY). We thank Chase Hintelmann, Prisha Patel, and Divya Raskonda for their assistance with data collection.

## Author contributions

MPB, GMN, and JDY performed the experiments; MPB, XZ, AC, KZ, and JDY analyzed the data; TMM and JDY designed the experiments; MPB, TMM, JDY wrote the paper.

## Data availability statement

All study data and analysis code can be found at https://rutgers.box.com/v/Berns-et-al-NIHL-Auditory.

