## Supplemental Figures for "Auditory decision-making deficits after permanent noise-induced hearing loss"

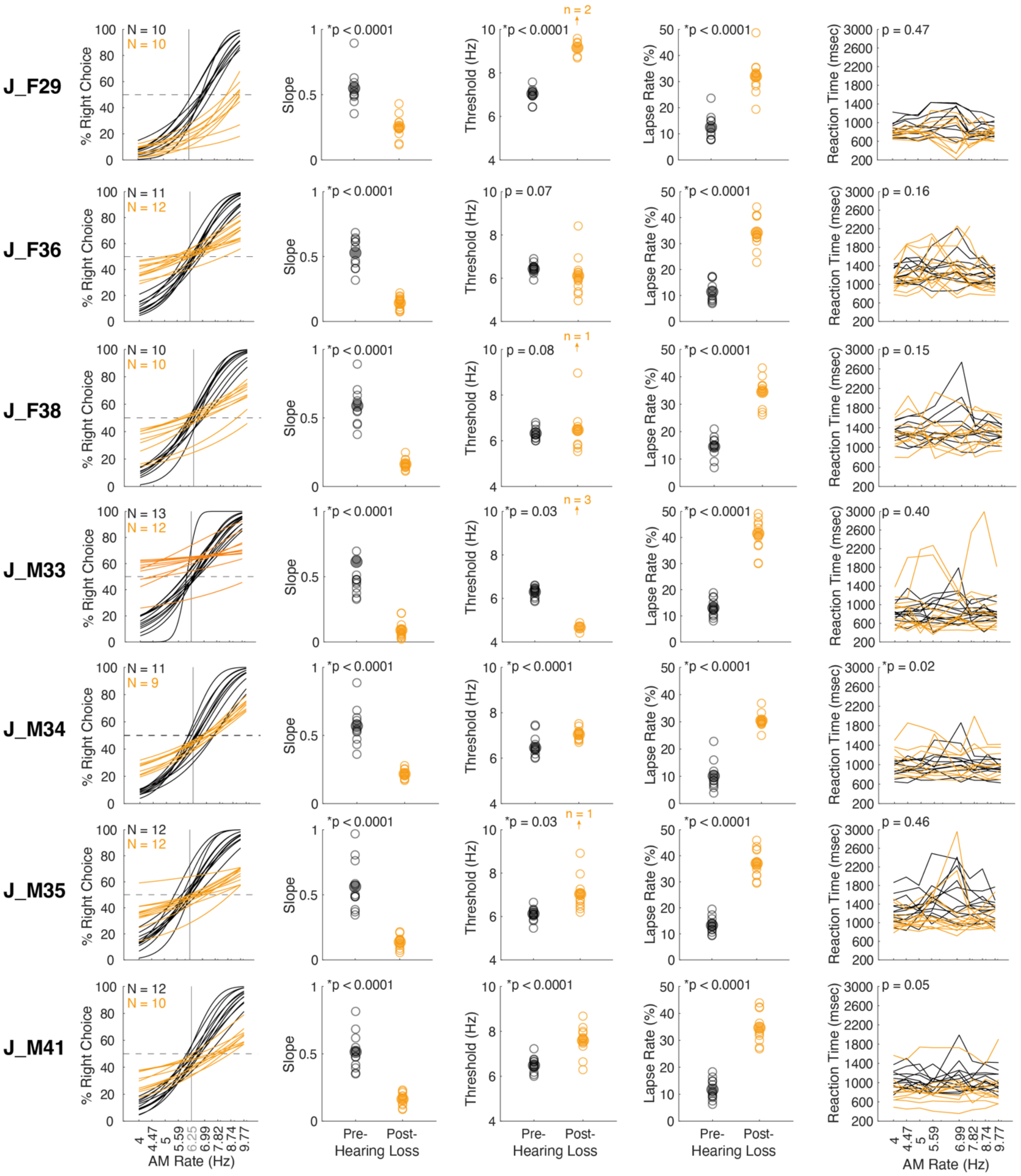


**Supplementary Figure 1**

Psychometric performance for all tested animals. For each animal (row), we plot individual psychometric functions, corresponding slope, threshold, lapse rate, and reaction times for correct trials from test sessions across pre- (black) and post-NIHL conditions. Asterisks denote statistically significant differences within each subject (two-way t-test).

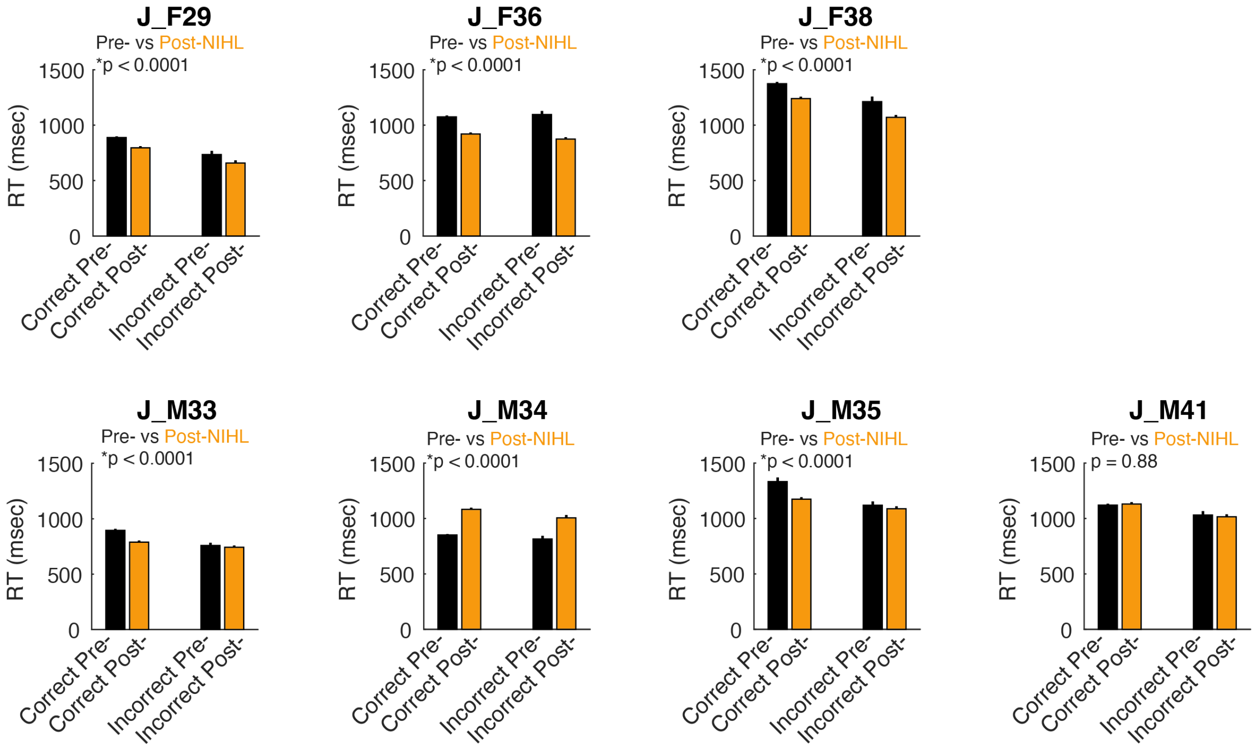


**Supplementary Figure 2**

NIHL can impact task performance reaction times (RTs). For each animal, we plot mean ± SEM RTs for different trial outcomes (Correct and Incorrect) across pre- (black) versus post-NIHL (orange) conditions. Asterisks denote a statistically significant differences of RTs between trial type and hearing status within each subject (two-way mixed model ANOVA).
